# Single influenza A viruses induce nanoscale cellular reprogramming at the virus-cell interface

**DOI:** 10.1101/2024.07.13.603385

**Authors:** Lukas Broich, Hannah Wullenkord, Maria Kaukab Osman, Yang Fu, Peter Reuther, Mark Brönstrup, Christian Sieben

**Affiliations:** Nanoscale Infection Biology Group, Helmholtz Centre for Infection Research, Braunschweig, Germany; Department of Chemical Biology, Helmholtz Centre for Infection Research, Braunschweig, Germany; Institute of Virology, Medical Center - University of Freiburg, Freiburg, Germany; Faculty of Medicine, University of Freiburg, Freiburg, Germany; Spemann Graduate School of Biology and Medicine, University of Freiburg, Freiburg, Germany; Institute of Genetics, Technische Universität Braunschweig, Braunschweig, Germany

**Author notes:** correspondence should be addressed to: Christian Sieben, Helmholtz Centre for Infection Research Nanoscale Infection Biology Group Inhoffenstr. 7, 38124 Braunschweig Germany.

**Keywords:** influenza, virus entry, endocytosis, single-molecule tracking, super-resolution microscopy

## Abstract

Viruses, as nanoscale entities with limited proteomes, must efficiently infect target cells using their available resources. During infection, individual virions induce specific cellular signaling within the virus-cell interface, a nanoscale patch of the plasma membrane in contact with the virus. However, virus-induced receptor recruitment and cellular activation are transient processes that occur within minutes and at the nanoscale level. Hence, the temporal and spatial kinetics of such early events often remain poorly understood due to technical limitations. To address this challenge, we developed a novel protocol to covalently immobilize unmodified influenza A viruses on glass surfaces before exposing them to live epithelial cells. This approach extends the observation time for virus-plasma membrane interaction while preserving the viruses’ native state for uncompromised cell interaction. Using single-molecule super-resolution microscopy, we investigated virus-receptor interaction showing that viral receptors are not immobilized by the virus but rather slowed down, which leads to a specific local receptor accumulation and turnover. We further followed the dynamics of clathrin-mediated endocytosis at the single-virus level and demonstrate the recruitment of adaptor protein 2 (AP-2), previously thought to be uninvolved in influenza A virus infection. Finally, we examined the nanoscale organization of the actin cytoskeleton at the virus-binding site, showing a local and dynamic response of the cellular actin cortex to the infecting virus. Our findings provide novel insights into the fundamental process of virus-cell interaction and demonstrate the versatility and potential impact of our approach on virus-cell biology.

## Introduction

Influenza A viruses (IAVs) are a major human health threat that cause annual epidemics and sporadic pandemics^1^. IAVs are enveloped particles, and their membrane harbors the two viral spike proteins hemagglutinin (HA) and neuraminidase (NA). To initiate viral replication and ensure the production of progeny virions, IAVs need to infect susceptible host cells. The infection process begins with binding of the viral HA to sialylated glycan attachment factors (AF) on the host cell plasma membrane^2^. While this initial interaction is critical for IAV host tropism^1^, it is of inherently low affinity^3^. We could recently show that IAVs overcome this limitation by forming multivalent contacts through binding of AF nanoclusters within the plasma membrane^4^. However, subsequent steps of the infection process remained elusive at the single-virus level.

Following AF binding, IAVs enter cells by clathrin-mediated endocytosis (CME) or macropinocytosis^5,6^. For CME-based cell entry, IAVs were shown to use pre-existing clathrin-coated pits or induced the *de novo* formation of new endocytosis sites suggesting a specific signaling function into the cell^7^. Since sialylated AFs do not possess any signaling capacity, functional receptor proteins were hypothesized early on and more recently a number of receptor candidates have been proposed to be involved in IAV cell entry^8–12^. Among them are tyrosine kinases such as the epidermal growth factor receptor (EGFR) or c-Met^11^. siRNA-mediated knock-down of EGFR in susceptible cell lines could reduce the infection rate of human H1N1 IAV strain A/Puerto Rico/8/34 (PR8) by about 60%^11^. Co-localization on the plasma membrane could also be shown, suggesting a direct interaction between IAV and EGFR^4,11^. We have recently performed single-virus tracking experiments to better understand how IAVs engage with functional EGFR nanodomains. Our results suggested that IAVs explore the plasma membrane in a SA-dependent way, thereby eventually reaching plasma membrane nanodomains that harbor both, sialylated AFs and functional EGFRs, which then become activated leading to downstream signal transduction and induction of CME^4^. However, the spatial and temporal visualization of this process at the single-virus level remained difficult due to the transient nature of the cell entry process and the small size of the IAV-cell interface.

In this work, we propose a new method to stabilize virus-cell interaction and enable a detailed view on the dynamics of the virus-cell interface including receptor recruitment and downstream CME induction. We developed a protocol to modify microscopy glass substrates for the immobilization of unmodified IAVs, thereby generating a stable viral interface that can then be brought in contact with live cells for microscopic investigation. With our method, we were able to prolong the plasma membrane-contact between virus and cells to investigate the interaction between IAVs and individual EGFR molecules, using advanced single-molecule localization microscopy (SMLM) and single-particle tracking. By determining the mobility of individual receptors, we could identify a significant decrease of receptor diffusion in proximity to IAV, indicating a direct and quantifiable interaction between receptor and virus. In subsequent experiments, we show the induction of CME as well as the dynamics and nanoscale response of the actin cytoskeleton at the virus binding site. Taken together, our approach allows, for the first time, a nanoscale view on IAV-induced cellular reprogramming at the virus-cell interface.

## Results

### Unmodified IAVs can be covalently linked to glass substrates before live cell attachment

To generate a stable virus-cell interface enabling high-resolution visualization of virus-cell interaction, we chemically modified glass coverslips for the covalent immobilization of native viruses and the attachment of live cells. The coverslips were coated with a silane-PEG_5000_-NHS linker that could further react with primary amines present on the viral spike proteins. To passivate the regions between the viruses and reduce the amount of NHS-linkers, silane-PEG_5000_-NHS was mixed with silane-PEG_200_ at a ratio of 1:1 (**Figure 1A**). We immobilized fluorescently-labeled IAV PR8 particles and quantified the number of attached particles per 500 µm^2^, which corresponds to the projected 2D surface area of an attached A549 human lung epithelial cell. The number of immobilized virus particles and their integrated intensity was determined by confocal microscopy (**Figure 1B**). In comparison to an unreactive PEG-passivated slide, we found a 2.5-fold higher number of viruses on the NHS-modified slide, with very few particles being attached to silane-PEG_200_ coated slides. We then investigated the intensity distribution of the immobilized IAV particles, which shows a single peak that overlaps with the intensity distribution of 100 nm fluorescent beads, recorded in a parallel experiment (**Figure 1C**). This indicates that virus particles did not aggregate during the labelling and immobilization procedure in spite of the pleomorphism of IAVs^13,14^. Together, this one-step silanization protocol provides a homogenously distributed viral attachment pattern on the microscopy glass slides.

**Figure 1.**
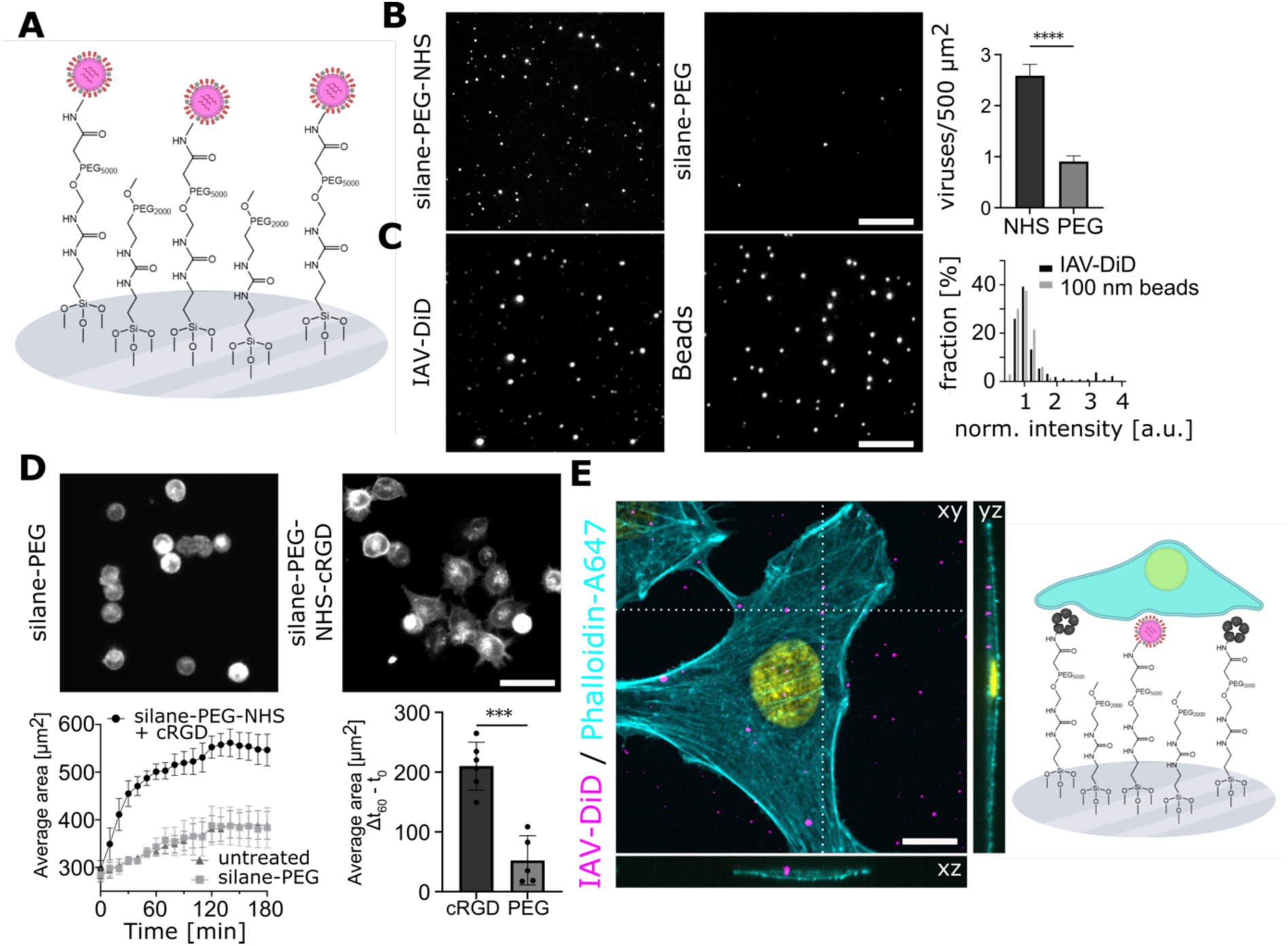
Unmodified IAVs can be covalently linked to glass substrates before live cell attachment. **A.** Schematic of the glass substrate modified with silane-PEG-NHS and silane-PEG. **B.** DiD-labeled IAV particles were incubated for 1h at RT on reactive glass slides coated with silane-PEG-NHS or silane-PEG. After subsequent washing, the number of bound viruses per area was quantified (right panel). Slides coated with PEG-NHS showed a statistically significant increase of bound viruses per 500 µm² compared to slides coated with only PEG (p < 0.0001, n = 200, bottom right). **C.** Fluorescent 100 nm beads and labeled IAVs were immobilized on silane-PEG-NHS-coated glass slides. The particles were counted and the fluorescence intensity was measured for each particle. Histograms of the normalized intensities were compared and show that both distributions scatter around the same mean value of ∼1 (t-test, p = 0.8359, n = 659) (right panel). However, the distribution of viral fluorescence intensities shows a significantly higher variance compared to the one obtained for fluorescent beads (F-test, p < 0.0001, n = 659). **D.** Fluorescently-labeled A549 cells were cultured on slides coated with silane-PEG or silane-PEG-NHS-cRGD. The average cell area was followed over time. A comparison of the growing cell area over the course of 3 hours shows that cells growing on the cRGD-coated slides adhere fastest (bottom left). After one hour, a significant increase in cell area was observed for cells growing on PEG-NHS-cRGD coated slides, compared to cells growing on slides coated with PEG alone (p = 0.0001, n = 11, bottom right). **E.** DiD-labelled IAVs were mixed with 0.5 mM cRGD and immobilized on a silane-PEG-NHS coated glass slide. After subsequent washing, A549 cells were cultivated on the slides for 16 hours. Afterwards, cells were fixed with 4% PFA and cellular actin was stained with phalloidin-A647. DNA was counterstained with DAPI (yellow). Confocal imaging confirmed that the cells are growing on top of the immobilized viruses (left). On the right site, a schematic of the final experimental setup is depicted, showing the cells growing on top of immobilized viruses and cRGD (not to scale). Scale bars: B, 10 µm; C, 10 µm; D, 50 µm; E, 10 µm.

The next step was to enable the attachment of live mammalian cells on top of the immobilized IAVs. To aid live cell attachment, we conjugated a small cyclic pentapeptide with an arginine-glycine-aspartate motif (Cyclo(-RGDfK), further referred to as cRGD), a subunit of the extracellular integrin-binding matrix protein fibronectin^15^, to the silane-PEG_5000_-NHS linker. We then analyzed the time course of cell adhesion using fluorescently-labeled A549 cells. Quantification of the cellular area over time showed that the cRGD coating indeed enhanced the speed of cell attachment with a saturation point of the growth curve already after 1 hour. Cell adhesion was slower and less pronounced on untreated and silane-PEG treated glass slides, as exemplified by the 30% smaller cellular area (**Figure 1D**). To adhere both, IAV and live A549 cells to the slide simultaneously for subsequent microscopic analyses, we mixed the labelled viruses with 0.5 mM cRGD solution and immobilized both at the same time onto the slides. We then cultured A549 on the immobilized viruses and used confocal microscopy to investigate the 3D organization of the sample. Following cell fixation and staining with Alexa488-conjugated phalloidin to visualize the cellular periphery, we observed that immobilized viruses are indeed located under the cells on the same slide (**Figure 1E**).

In summary, our glass modification protocol allows the immobilization of native unmodified virus particles and is compatible with the attachment of live mammalian cells. Our platform can now be used to investigate the formation and signaling function of the virus-cell interface using fluorescence microscopy.

### Immobilized IAVs recruit EGFR in a sialic acid-dependent way

To successfully infect a cell, viruses need to interact with plasma membrane receptors. While it is well described that IAVs bind host cells via sialic acid, the nature of the functional entry receptor remains less well understood. Previously, EGFR was suggested as a potential candidate receptor for IAV PR8^11^. To investigate the role of EGFR for the infection of host cells by IAV, we immobilized fluorescently-labeled IAV PR8 particles on glass surfaces and cultured A549 cells expressing endogenous levels of EGFR-GFP on top of these viruses. We then used live-cell total internal reflection (TIRF) microscopy to investigate the distribution of EGFR-GFP in proximity of immobilized viruses. We found an accumulation of the EGFR-GFP signal colocalizing with fluorescently-labelled IAV particles (example in **Figure 2A**). To test whether this accumulation was dependent on HA-sialic acid interactions, the cells were treated with 250 mU/ml sialidase for 1h prior to imaging. The accumulation of EGFR-GFP was not visible anymore, indicating a sialic-acid-dependent IAV-EGFR interaction (**Supplementary Figure 1**).

**Figure 2.**
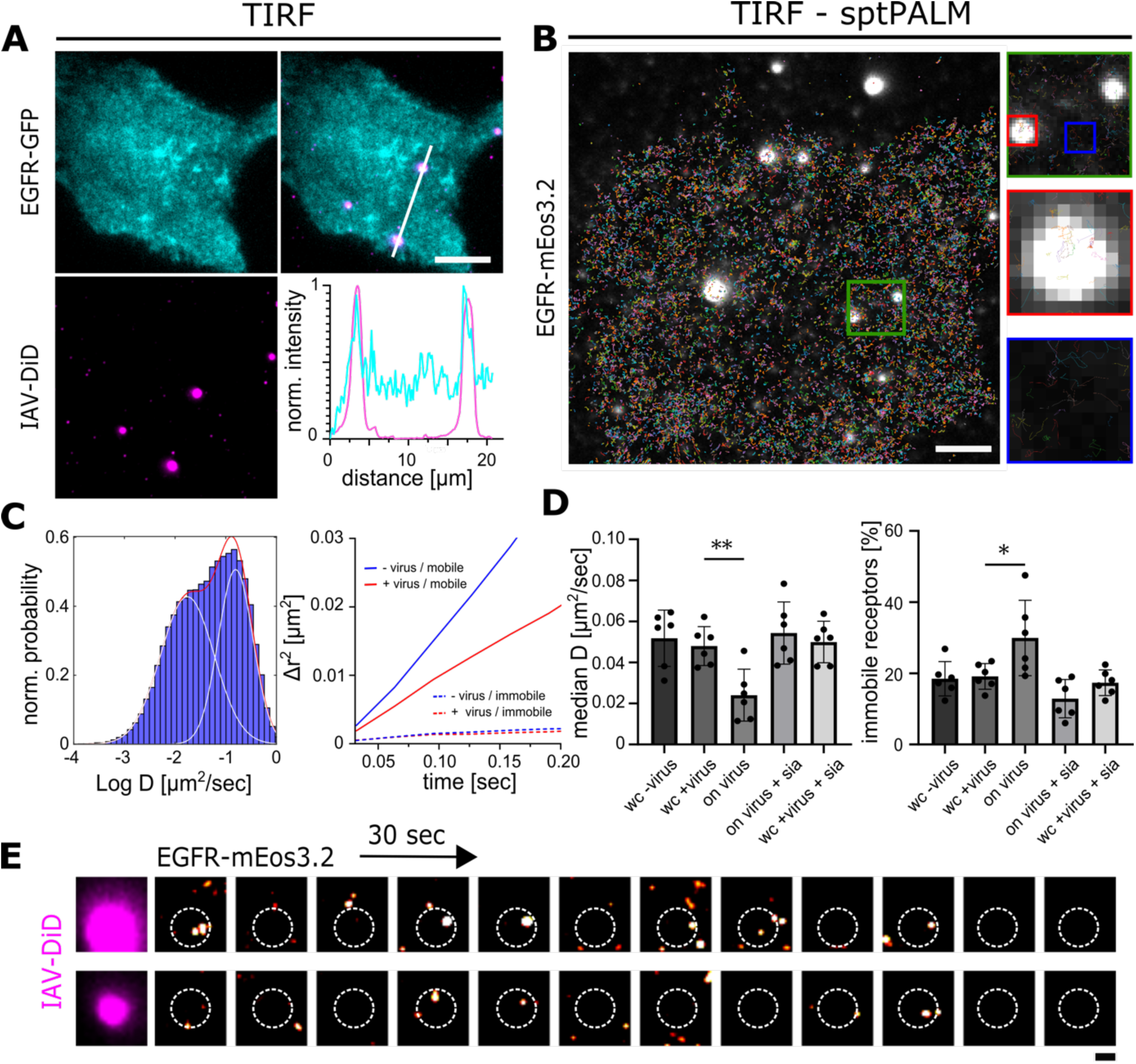
Immobilized IAVs recruit EGFR in a sialic acid-dependent way. **A**. IAVs were immobilized on reactive glass surfaces, and A549 cells expressing EGFR-GFP were cultivated on top. Live-cell TIRF microscopy reveals EGFR signal accumulation at the virus binding sites. The bottom right line plot shows the normalized intensities along the white line shown in the overlay picture above. **B.** A549 cells expressing EGFR-mEos3.2 were cultivated on top of fluorescently labeled IAVs. SptPALM imaging was applied and EGFR tracks were reconstructed. Plots of the tracks were superimposed on a still image of the underlying viruses. Tracks are color-coded to remain distinguishable. Further analysis was performed comparing tracks in viral proximity (red square) with tracks in membrane patches without virus (blue square). Zoom of the green region shown on the right. **C.** MSD analysis of EGFR trajectories taken from cells growing on immobilized viruses (whole cell) shows a bimodal distribution with a mobile and an immobile/confined population. **D.** Initial diffusion coefficients were calculated from the MSD curves of individual EGF receptors at different conditions (left panel). For each bar, the median initial diffusivities of EGFR from six experiments were used. In each experiment, an average of ∼10k receptors was tracked. The average median diffusivity of receptors in cells growing on reactive slides with and without viruses is not statistically different (whole cell (wc) ± virus). However, looking exclusively at receptors in an 800 x 800 nm square around immobilized viruses (on virus) shows a significantly reduced diffusion coefficient (p = 0.0040, n = 6). This effect is reversible by treating the cells with sialidase for 1h before the measurement (on virus + sia and wc +virus +sia). Fractions of immobile receptors were analyzed from sptPALM experiments (right panel). Receptors with an initial diffusion coefficient D_1-3_ < 0.01 µm²/s are considered immobile. In viral proximity, a statistically significant increase in immobile receptor fractions can be observed (p = 0.0398, n = 6 fields of view). **E.** EGFR-mEos3.2 localizations from a live-cell acquisition were rendered using 30 sec time binning resulting in 12 reconstructions across a 10 min acquisition. The position of the labelled IAV particle is shown on the left and indicated as a dotted circle on the PALM reconstructions. Recurrent appearance of EGFR clusters can be observed indicating dynamic exchanges of EGFR between the virus-interface and the remaining plasma membrane. Scale bars: A, 10 µm; B, 5 µm; E, 1µm.

Virus-receptor interaction occurs within the virus-cell interface, a nanoscale patch of the plasma membrane. To gain deeper insights into the dynamic interaction between IAVs and individual EGFR molecules, we used photoactivated localization microscopy (PALM), a super-resolution microscopy method that relies on the sparse photoswitching and subsequent localization of individual photoactivatable proteins^16,17^. For this purpose, A549 cells expressing EGFR tagged with mEos3.2, a PALM-compatible fluorophore^18^, were cultured on top of immobilized IAV particles. Individual live cells were imaged for 300 s and EGFR-mEos3.2 localizations determined using established single-molecule localization routines (see methods). Importantly, since we used live cells, EGFR molecules could be followed (tracked) over several frames by single-particle tracking PALM (sptPALM^19^), which allowed the localization of individual receptor molecules and the reconstruction of their movements with nanometer precision. On average about 10,000 receptor trajectories were reconstructed per cell. For each EGFR trajectory, the mean squared displacement (MSD) as well as the initial diffusion coefficient (D_1-3_) was calculated (see methods).

For cells grown on linker-modified slides without any immobilized viruses, we determined the median initial diffusion coefficient (D_1-3_) for EGFR to be 0.051 ± 0.013 µm²/s. Notably, similar values of approximately 0.05 µm²/s or 0.048 ± 0.065 µm²/s have been reported for EGFR in A431 cells, a comparable human epithelial cell line^8,9^. This indicates that the modified glass surface did not have a negative impact on the diffusion properties of EGFR.

To then investigate the effect of virus binding on the EGFR dynamics, we repeated the experiment with cells cultured on top of immobilized IAV particles. Median diffusivities were compared either at the whole cell (WC) level or for receptors found in 800 x 800 nm squares around immobilized viruses (**Figure 2B**, red square). We determined the initial diffusion coefficient (D_1-3_) of all EGF receptors to be 0.040 ± 0.009 µm²/s (**Figure 2C, D**). This is within one standard deviation from the measurements without virus (**Figure 2D**, **Supplementary Figure 2**). We thus conclude that the mere presence of the virus does not appear to alter the overall receptor diffusivity at the whole-cell level. MSD vs. lag time plots allow to discriminate different particle movement patterns by analyzing the shape of the resulting curve^20^. Comparing MSD vs. lag time plots of receptors in viral proximity with receptors in other membrane patches (**Figure 2B**, red/blue squares) showed a distinct diffusive behavior for both groups (**Figure 2C**, right panel) indicating slower EGFR diffusion in viral proximity (+/- virus mobile). Indeed, when we specifically looked at receptors within membrane patches located around the viruses, we observed a significant reduction in diffusivity to 0.024 ± 0.012 µm²/s (**Figure 2D**, on virus, p = 0.0040). To again test the effect of sialic acid on the reduction of IAV-mediated EGFR slowdown, we treated the cells with sialidase before the measurement. Sialic acid digestion reversed this effect, resulting in a diffusion coefficient of 0.054 ± 0.015 µm²/s for receptors in viral proximity (**Figure 2D**, on virus + sia). We further compared the fraction of slowed receptors between the different samples, considering receptors with D_1-3_ < 0.01µm²/s as slowed down^10,11^. A significantly higher fraction of slowed receptors was observed in proximity of the viruses (**Figure 2D**, right panel, p = 0.0398).

Finally, we wanted to investigate the structural organization of EGFR within the virus-cell interface. Interestingly, while we observed EGFR clusters as reported previously^4^, they appeared only transiently over time, indicating exchange of receptors from the plasma membrane reservoir (**Figure 2E**). Due to the shortness of sptPALM trajectories, EGFR movement in and out of the virus-cell interface could only rarely be observed. We thus used EGFR-GFP-expressing A549 cells and indeed observed a long-term exchange (minutes) of EGFR molecules within the virus-cell interface (**Supplementary Figure 3**).

In summary, these results suggest that immobilized IAVs form a functional virus-cell interface capable of binding EGFR, and this interaction depends on sialic acid. The virus-cell interface is not a static platform where receptors become immobilized; instead, the receptors are slowed down, allowing a characteristic enrichment (**Supplementary Figure 4**) and dynamic exchange of receptors from the larger plasma membrane reservoir.

### Immobilized IAVs recruit proteins involved in clathrin-mediated endocytosis

After we could observe and quantify the interaction between IAV and host cellular EGFR, we wanted to test if the interaction induced downstream cellular signaling. The subsequent major event of IAV infection would be the induction of endocytosis, a crucial step leading to the internalization of viral particles into the host cell cytoplasm. For non-immobilized IAVs, CME has been shown to be the main endocytosis pathway. Depending on the IAV strain, the majority of viruses (∼60%) is internalized by CME, while the remaining fraction uses clathrin-independent pathways^7,21^. Further research suggested that CME-based IAV entry is dependent on the adaptor protein epsin-1^22^.

To further assess the functionality of our assay, we cultured A549 cells expressing epsin1-GFP on top of immobilized IAV particles (**Figure 3A**). The cells were imaged using TIRF microscopy under live cell conditions. We found that epsin-1-GFP displayed discrete point-like structures and that 38 ± 8% (n = 49) of immobilized IAVs colocalized with epsin1 similar to the previously published value (∼50%) for apical IAV infection^22^.

**Figure 3.**
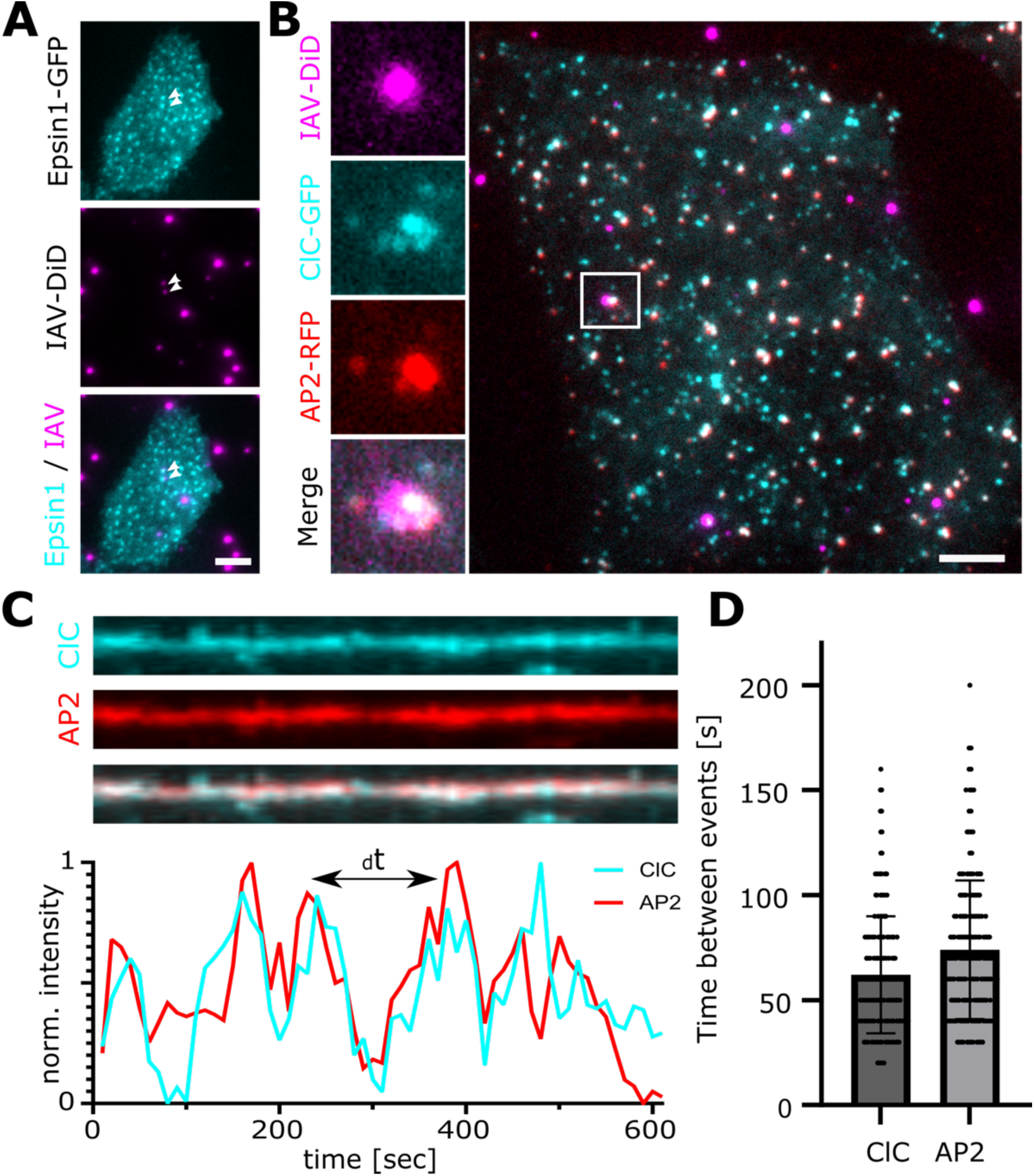
Immobilized IAVs recruit proteins involved in clathrin-mediated endocytosis. **A.** A549 cells were transfected with epsin1-GFP and cultured on top of immobilized IAVs. In TIRF microscopy, a colocalization of epsin1-GFP was observed for 38 ± 8% of viruses (n = 49). **B.** IAVs were immobilized, and MDA-MB-231 cells co-expressing endogenous levels of AP2-RFP and ClC-GFP were cultured on top. TIRF imaging revealed colocalization of AP2 and ClC with 63.33 ± 12.02% (n = 45) of viruses. Zoom of the white boxed area shown on the left. **C.** Cell-virus interactions from B have been captured over the course of 10 minutes with dual-cam live cell TIRF microscopy. The frequency of AP2-RFP and ClC-GFP localizations at the virus binding site was quantified from the extracted time traces, as shown in the respective kymograph. **D.** The ClC-GFP signal peaked on average every 62.1 ± 27.93 sec at the virus binding sites. The AP2-RFP signal peaked on average every 73.94 ± 32.95 s (n = 28). Scale bars: A, 5 µm; B, 5 µm.

EGFR was shown to associate with different classes of clathrin-coated pits (CCPs), which are discriminated by the presence of adaptor protein 2 (AP2), where AP2-negative CCPs would rely on other endocytic adaptor proteins such as eps15 and epsin-1^23^. AP2 is a major constituent and pioneering factor of CCPs at the plasma membrane. To test if immobilized IAVs also recruit AP2 to the plasma membrane, we used cells stably expressing endogenous levels of AP2-RFP together with clathrin light chain A (ClC) fused to GFP. These cells were then cultured on top of immobilized IAVs and observed by live-cell TIRF imaging. We found that 63.33 ± 12.02% (n = 45) of immobilized viruses colocalized with both AP2 and ClC (**Figure 3B**), confirming that IAVs indeed recruit AP2 to the plasma membrane. Interestingly, when we monitored the signal intensity over a period of 10 minutes, we detected fluctuations in the fluorescence signals for both ClC-GFP and AP2-RFP, indicating a recurring interaction between the virus and the cellular CME machinery. We identified signal peaks, quantified the average time between two intensity peaks, and found that ClC-GFP peaks had a mean interval of 62.10 ± 27.93 s, while AP2-RFP peaks could be observed every 73.94 ± 32.95 s (**Figure 3D**, n = 28). Similar recruitment dynamics have been shown before for reoviruses immobilized by click-chemistry^24^, which are in a similar size range as IAV particles.

From these experiments, we conclude that immobilized IAVs are in a continuous interaction with their host cells, with repeated formation of clathrin-coated pits at the viral binding sites. We document recruitment of epsin-1 as shown before^22^ but surprisingly also AP2, which was previously suggested to be dispensable for IAV infection^25^.

### Immobilized IAVs lead to local redistribution of the cellular actin cytoskeleton

The dense cortical F-actin network beneath the plasma membrane plays a central role in many cellular processes including CME. During CME, polymerization of the globular G-actin into the filamentous F-actin was suggested to support membrane deformation and scission^26^. While IAV cell entry was shown to depend on the actin cytoskeleton only in polarized cells^27^, a recent phosphoproteomic study revealed a fast cellular re-programming of actin regulatory proteins upon IAV-cell attachment^28^. We thus hypothesized that IAV-cell binding induces a local F-actin reorganization.

To investigate the F-actin organization precisely at the virus-binding site, we stained the cellular F-actin using Alexa647-conjugated phalloidin in A549 cells cultured on top of immobilized IAV particles (**Figure 4A**). We imaged the cells in TIRF to visualize F-actin structures close to the plasma membrane and found that about 15% of IAVs show a distinct accumulation of F-actin (**Figure 4A, Supplementary Figure 4**). To further investigate the nanoscale organization of the F-actin accumulation in proximity of cell-bound IAVs, we performed stochastic optical reconstruction microscopy (STORM). STORM is a super-resolution microscopy method that relies on the chemically-induced photoswitching of organic dye molecules, which allows their localization with nanometer precision^29^. STORM reconstructions revealed the complex and diverse nanoscale organization of the F-actin network within the TIRF field of adherent A549 cells (**Figure 4B**). While we could detect many larger filamentous structures with diameters around 89 ± 21 nm (FWHM of 20 filaments), immobilized IAVs seem to overlap with distinct structures that we then classify based on their appearing structural organization as 1) arc or ring-like structures, 2) large clusters (blobs) or 3) areas without any distinct F-actin accumulation (holes) (**Figure 4B**, white zoom boxes). With respect to previous reports on the nanoscale organization of F-actin during CME^30,31^, we hypothesized that the observed structures (1-3) are intermediates of the same process. We thus performed live-cell imaging of F-actin at the virus-binding site. To this end, we cultured A549 cells expressing LifeAct-GFP on top of immobilized IAV particles (**Figure 4C, Supplementary Video 1**). The cells were imaged using TIRF microscopy under live-cell conditions. We observed a similar diversity of structures as observed after phalloidin staining, indicating that F-actin was efficiency labelled in both cases^32^. When following the LifeAct-GFP signal in proximity of individual viruses (**Supplementary Video 1**), we could indeed observe fluctuations with transitions between different structures that resemble the patterns (1-3) observed by STORM. Snapshots of the LifeAct-GFP fluctuations around one virus particle (**Figure 4C, insert**) are shown in **Figure 4D**. Line plot measurements (**Figure 4C, insert, white line**) disclosed the different F-actin patterns transitioning from a hole to a ring and into a blob that overlaps with the viral signal before turning back into a ring-like organization.

**Figure 4.**
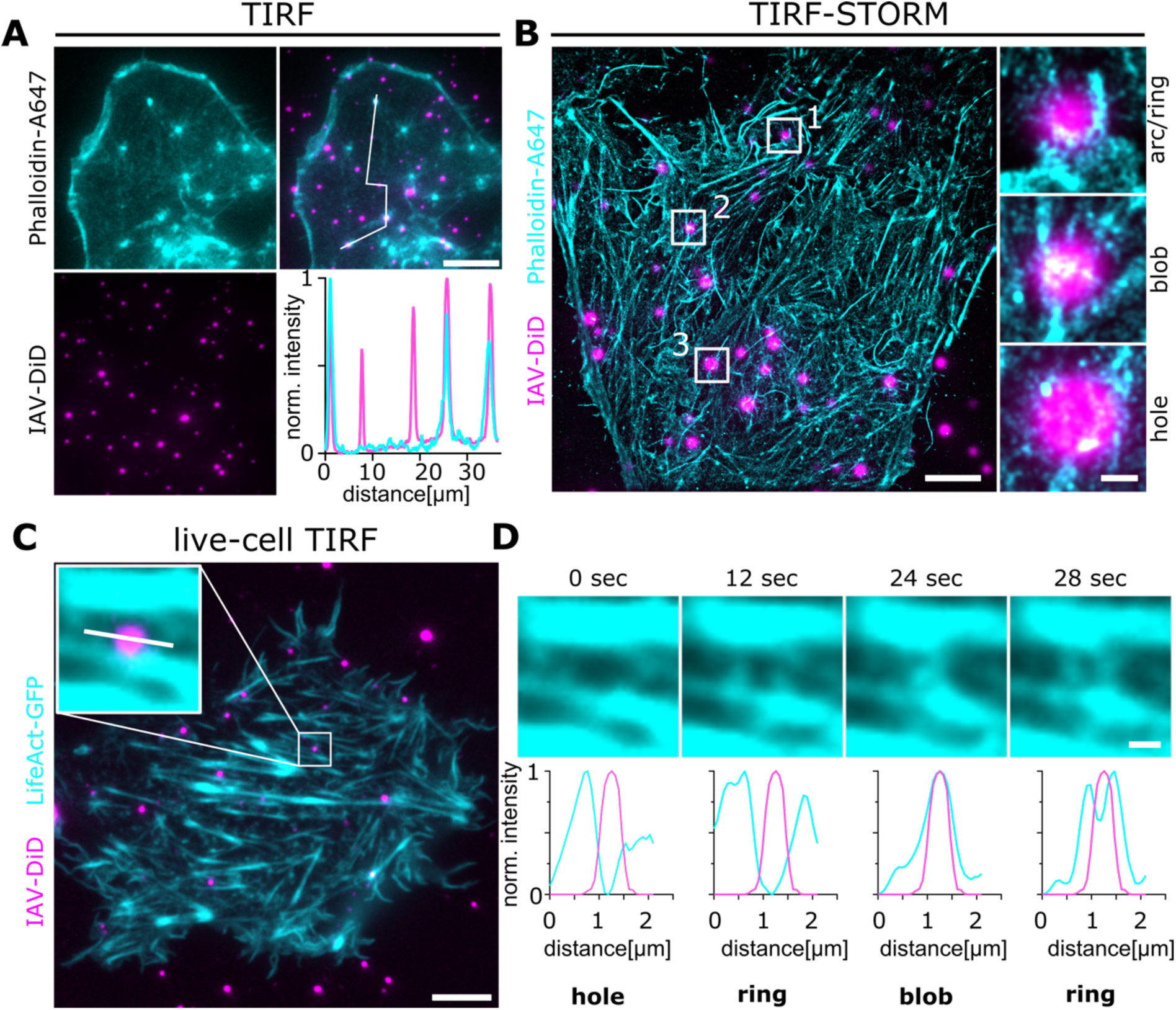
Local re-organization of the actin cytoskeleton at the virus-cell interface. **A**. DiD-labelled IAVs were immobilized on reactive glass surfaces and A549 cells were cultivated on top. The cells were fixed and the actin cytoskeleton was labelled with Alexa647-conjugated Phalloidin. The sample was imaged using TIRF to visualize actin structures close to the plasma membrane. We observed that some viruses show a distinct accumulation of the phalloidin signal as visualized by a line plot measurement (white line, upper right panel). **B.** To further investigate the nanoscale organization of F-actin at the virus binding site, we performed STORM using the same samples as shown in **A**. We identified a diverse F-actin organization with many filamentous structures as well as some smaller local accumulations. Closer inspection of the A647-phalloidin signal in proximity of immobilized viruses revealed different structural patterns that we classify as arcs/rings, holes or local amorphous accumulations (blobs) (**B**, right side). **C**. We performed live-cell TIRF imaging of A549 cells expressing Lifeact-GFP and cultivated on top of immobilized DiD-labelled IAVs. While we observed a similar diverse structural organization of Lifeact-GFP, we could now follow its signal at the viral periphery over time (**D**). **D** shows snapshots of a 5 min movie of the cell shown in **C** zoomed at the viral particle shown in the boxed inset. The four snapshots and the corresponding line profiles below show the structural transitions of the local Lifeact-GFP signal. Scale bars: A, 10 µm; B, 5 µm; B, zoom boxes, 500 nm; C, 10 µm; D, 1 µm.

From these experiments, we conclude that immobilized IAVs induce local F-actin re-organization that can be imaged in live cells or investigated by STORM. Our findings highlight the dynamics of the local F-actin cytoskeleton in response to an attached virus and open many possibilities towards exploring the functional implications of the observed structures.

## Discussion

Virus-cell entry is a dynamic process that involves the recruitment of cellular receptor proteins and their activation, which is followed by specific downstream signaling events that eventually lead to virus endocytosis. Despite the relevance of this process for virus infection, its transient nature and the small scale of the virus-cell interface have made it difficult to study at high spatial and temporal resolution. Other strategies to immobilize virus particles have been developed before but require genetic or chemical modification of viral particles^24,33^. These assays are time- and resource-intensive and may also be impractical for certain viruses as the modification of the spike proteins might affect their ability to successfully infect a cell. We have developed a universal method to immobilize native virus particles on glass slides compatible with subsequent cultivation of live mammalian host cells. Our method allows to investigate the early interaction between an infecting virus and its host cell at the single-virus level. Importantly, the technique has the capability to measure the interaction between a host cell and multiple viruses at the same time and is compatible with a variety of different microscopy methods, including single-molecule super-resolution microscopy. We have demonstrated the versatility of our approach by studying IAV-receptor interaction, CME induction and local F-actin reorganization, key steps involved in IAV endocytosis.

As a model system, we chose the human IAV strain A/Puerto Rico/8/34 (PR8), which was shown before to activate and use EGFR for cell entry^11^. We could previously show that IAVs use plasma membrane nanodomains for multivalent cell attachment and subsequent activation of EGFR^4^. Here we applied sptPALM of EGFR-mEos3.2 to investigate the dynamics of IAV-EGFR interaction in live cells. We found that IAV binding slows down EGFR within the virus-cell interface and that this reduced mobility is sialic acid-dependent. We then reasoned that an active virus-cell interface should be able to recruit downstream proteins, such as those involved in CME. We could indeed observe recruitment of epsin-1 to the virus binding site as previously shown for IAV strain X-31^22^. Since IAV PR8 and X-31 both activate EGFR^34^, which itself was shown to use multiple CME pathways, we further tested the recruitment of AP2 and clathrin to the virus binding site. Although AP2 was previously suggested not to be involved in IAV-cell entry^25^, we found recurring recruitment of both proteins with frequencies similar to those observed before for reoviruses^24^. From these experiments, we conclude that immobilized viruses induce CME and that this process involves both epsin-1 and AP2-dependent pathways. While we show a specific sialic acid-dependent slowdown of receptor proteins, we cannot exclude curvature-dependent CME induction independent of receptor activation, as previously shown for reoviruses and synthetic surfaces^24,35^. Because most IAVs use multiple entry receptors^36^, our assay could be applied for more detailed studies to elucidate the contribution of specific receptor proteins in IAV-cell entry. In a parallel study, we have used the inverted infection approach to investigate the interaction between H18 IAVs and MHCII receptors. We found a more pronounced receptor slowdown as compared to the PR8-EGFR case, which was accompanied by a strong receptor clustering^37^. Lastly, we use our assay to probe the local response of the actin cytoskeleton to virus-cell binding. We observed different structural arrangements (ring, hole, blobs) that are consistent with similar structures observed in live-cell imaging. The small fraction of viruses (∼15%) showing a distinct colocalization with Lifeact-GFP could be a result of the transient nature of the observed F-actin fluctuations. Also, while CME in yeast is strictly actin-dependent^38^, not all CME sites in mammalian cells seem to require actin accumulation^39^, here presumably to counteract membrane tension^27,40^. We thus conclude that the observed structures are intermediates of the same process. Since viruses cannot be internalized, the actin fluctuations are recurring as we also show for the recruitment of CME-associated proteins.

In summary, we believe that our method will facilitate research on processes occurring during early viral infection. Moreover, because the method can easily be applied to other cargo particles such as synthetic amine-functionalized nanoparticles, virus surrogates or bacteria, it may find utility for multiple areas of infection research on the single-cell level.

## Acknowledgements

C.S. would like to thank Jennifer Fricke for her excellent technical assistance. This work was supported by the Helmholtz Association (VH-NG-1526 to C.S.) and a PhD fellowship of the China Scholarship Council (CSC to Y.F.). C.S. and M.B. acknowledge support through the cooperativity and creativity project call (CCC, project 8) of HZI. M.K.O. is a member of the Spemann Graduate School of biology and medicine (SGBM). P.R. is supported by the Hans A. Krebs Medical Scientist Programme of the Medical Faculty of the University of Freiburg.

## Materials and Methods

### Cell lines and culture

A549 wild-type cells (A549 WT, ATCC: CCL-185) and A549 cells endogenously expressing GFP-tagged EGFR (Sigma-Aldrich, CLL1141) were cultured and maintained in DMEM (Gibco, 10566016) supplemented with 10% FCS (Capricorn, FBS-LE-12A). A549 WT cells were modified by retroviral transduction (see c) to generate a stable cell line overexpressing EGFR-mEos3.2. To keep the selection pressure on these cells, 1.25 µg/ml puromycin (Gibco, A1113803) was added to the culture medium. MDA-MB-231 cells (provided by David Drubin, UC Berkeley) endogenously expressing ClC-GFP and AP2-RFP were cultivated in DMEM/F-12 (Gibco, 10565018) medium supplemented with 10% FCS. All cells were maintained in a humidified incubator at 37°C and 5% CO_2_ and passaged every 3 to 4 days. For passaging or seeding cells for an experiment, cells were detached using 0.5% Trypsin/EDTA (Gibco, 25300054), counted and resuspended in fresh growth medium at the desired concentration. Prior to all measurements cells were starved in FCS and phenol red free medium for 1h (Gibco, 31053028). For sialidase treatment, cells were starved in the same medium supplemented with 250 mU/ml sialidase (*Clostridium perfringens,* Roche, 11585886001).

### Influenza virus production

The viruses used for these experiments were produced in specific pathogen free eggs from ValoBiomedia. Eggs were incubated for 10 days at 37°C and 50% – 70% humidity and rotated regularly. Influenza A/Puerto Rico/8/34; H1N1 (kindly provided by Klaus Schughart, HZI) was diluted 1:1000 in sterile PBS. Egg surfaces were disinfected (100 mg/ml Povidone-iodine) and punctuated. 200 µl virus suspension was injected into the eggs and the injection site was sealed with glue. Subsequently, eggs were incubated for 48h at 37°C and 50% – 70% humidity, without rotation. After the incubation period, eggs were stored at 4°C overnight. Eggs were opened and allantoic fluid was aspirated and collected. Virus content of each sample was individually assessed in a hemagglutination assay (Chicken RBC, Fiebig Animal blood Products). Positive samples were pooled together, centrifuged at 2000g at 4°C for 30 minutes, aliquoted and frozen at -70 °C.

### Lentivirus production and transduction

To generate an EGFR-mEos3.2 overexpressing cell line, A549 WT cells were genetically modified by retroviral transduction. Retroviruses were produced in HEK-293T cells (ATCC: CRL-3216) cultured in a 6-well plate. When cells were at ∼70% confluency, they were co-transfected with a lentiviral packaging plasmid (psPAX2, Addgene #12260), a lentiviral envelope plasmid (pMD2.G, Addgene #12259) and a lentiviral transfer plasmid, using Lipofectamine 2000. The transfer plasmid encodes EGFR-mEos3.2 as well as a puromycin resistance (custom construct by Genescript, pLVX-M-puro backbone, Addgene #125839). Two days post-transfection the cell culture supernatant was harvested and cleared of cellular debris by centrifuging for 5min at 1300g. The supernatant was transferred to A549 cells growing in a 6-well plate in culture medium at ∼30% confluency. To facilitate viral infection the medium was supplemented with 20µg/ml polybrene (Sigma-Aldrich, TR-1003-G). Two days post-transduction the infection medium was removed and cells were cultured in growth medium supplemented with 1.25µg/ml puromycin for selection.

### Glass slide preparation

To clean and activate the glass coverslips (round 25 mm, No. 1.5, Epredia and 24 mm x 60 mm, No. 1.5, Marienfeld Superior), they were submerged in a freshly prepared piranha solution (5:1 H_2_SO_4_/H_2_O_2_) for 1 hour. Subsequently, the slides were washed twice with deionized water, dried using a nitrogen stream, and stored for up to 1 month at room temperature in a desiccator under a nitrogen atmosphere. To modify the activated surface, slides were treated with different ratios of silane-PEG_2000_/silane-PEG_5000_-NHS (tebu-bio, Offenbach, Germany) dissolved in dry DMSO for 45 min at RT. As negative control, an unreactive surface consisting of only silane-PEG_2000_ was used. Slides were washed three times in dry DMSO and twice in HEPES (0.1 M, pH 8) immediately before the subsequent reaction to avoid hydrolysis of the unstable NHS ester. As an alternative source, we also tested and performed experiments on commercially available reactive glass slides (3D epoxy glass slides, PolyAn, Berlin).

### Virus labeling and sample chamber preparation

To fluorescently label IAVs, lipid-dye conjugates (DiO, DiI and DiD, Invitrogen, V22889) were incorporated into the viral membrane. IAVs were diluted to a protein concentration of 1mg/ml in a final volume of 50 µl PBS. 1 µl of 10 mM lipid-dye solution was added and the suspension was shortly vortexed, spun down and incubated in the dark at RT for at least 1 hour. To separate unbound dye from labeled viruses, the suspension was fractionated by gel filtration (NAP-5, Cytiva 17-0853-01). Absorbance of the collected fractions was measured at 280 nm together with the respective dye absorbance wavelength (488 nm, 561 nm or 647 nm) to determine protein and dye concentrations (Nanodrop One, Thermofisher). Fractions that showed a peak at both wavelengths were collected and pooled together. To remove bigger aggregates of viruses, the viruses were mixed with 400 ml PBS and sterile filtered over a 0.22 µm filter right before immobilization. Sterile filtered viruses were diluted 1:150 in HEPES (0.1 M, pH 8) buffer, homogenized and added to the slide. Nanobeads (TetraSpeck, Invitrogen, T7279), were diluted 1:100 in HEPES before immobilization. Slides were kept at 4°C for 16h or for 1 h at RT. To measure multiple conditions on one slide, sticky-slide 8 well high chambers (Ibidi, 80828) were glued on to the slides. After incubation, the virus suspension was removed and the slides were washed three times with PBS, prior to seeding cells. Per chamber, 25,000 cells were seeded for all experiments.

### Cellular adherence measurement

Slides were prepared as described above. 0.05 mM cRGD (Cyclo(-RGDfK, MedChemExpress, HY-P0023) in HEPES (0.1 M, pH 8) was added and incubated for 1h to covalently couple cRGD to the NHS moiety on the linker. Slides were washed 3x in deionized water and dried in a nitrogen stream. Hoechst/DiO stained A549 cells (density: 1.25 x 10^5^ cells/ml) were seeded on top of different slides (untreated, 20 mg/ml silane-PEG_2000_, 20 mg/ml silane-PEG_5000_-NHS with cRGD), and cell adherence (area/time) was observed over the course of 3h. Attachment was monitored using a Keyence BZ-X800 fluorescence microscope with a stage top incubation chamber (Tokai Hit STR) and images were taken every 10 minutes. Adherence of the cells was analyzed using CellProfiler^41^.

### SMLM and image processing

SMLM was performed on a Nikon Ti Eclipse N-STORM microscope equipped with four laser lines at 405 nm (Coherent), 488 nm (Sapphire, Coherent), 561 nm (Sapphire, Coherent) and 646 (MPB Communications). The sample was observed through a Nikon Apo TIRF 100x Oil DIC N2 NA 1.49 objective and emitted light was detected on an Andor iXon3 DU-897 EMCCD camera (Oxford Instruments). Emitted light was directed onto the camera using a dichroic mirror and filtered through a Bright Line HC 609/64 (PALM) emission filter. We typically took between 10k (PALM) and 40k (STORM) frames at 30 ms integration time using the NIS elements software. Single emitter positions were localized using the software DECODE^42^. Separate models were trained for STORM and sptPALM acquisitions. To obtain the microscope-specific parameters needed to train a DECODE model, z-stacks of 100 nm fluorescent beads (TetraSpeck, Life Technologies, T7279) were acquired with a step size of 10 nm. Z-stacks were analyzed using SMAP^21^ to determine the necessary fitting parameters. These values were used as initial parameters to train the DECODE models. Each model was trained until it reached the maximum number of epochs or the loss-function did not reasonably improve anymore. Localizations were further processed using custom MatLab (MathWorks) or Python routines. For STORM, the localization were drift corrected and reconstructed using Thunderstorm^17^.

### sptPALM

A549 cells expressing EGFR-mEos3.2 were seeded on top of the immobilized viruses and incubated over night until sufficiently attached to the glass surface. Prior to imaging, the growth medium was removed and cells were starved for 1h in infection medium (DMEM, 0.2% BSA). For sialidase treatment, 250 mU/ml sialidase were added to the infection medium. Images were acquired in HEPES buffered DMEM (Gibco, 21063029) using a Nikon Ti Eclipse N-STORM microscope. For each measurement, 10,000 frames were acquired. A standardized procedure was use to filter the resulting single emitter positions: The initial 10% of frames (1000 frames) were discarded for all measurements as well as the 5% of localizations with highest uncertainties in x and y direction. Additionally, the 2.5% dimmest and brightest localizations were discarded. From the remaining localizations, trajectories were reconstructed with the use of the TrackPy^18^ python library. Localizations were linked using a KDTree algorithm with a maximum search range of 0.8 pixel and a zero-frame-gap memory. The adaptive stop for solving oversized subnets was set to 0.1 with a step size multiplicator of 0.9. For further analysis, only tracks with a minimum length of 10 frames were considered. Ensemble drift xy(t) was calculated and subtracted from the tracks. Mean squared displacement (MSD, equation 1) was determined and plotted individually for all molecules. For all tracks, the initial diffusion coefficients (D_1-3_) were calculated from the initial slope of the first three data pairs of individual MSD plots, by applying a linear fit in log space (Equation 2)^19,20^. MSD and diffusion coefficients for virus spots were calculated within an 800×800nm square around an immobilized virus. Receptors with D_1-3_ < 0.01µm²/s were considered immobile.^10,11^

Equation 1:

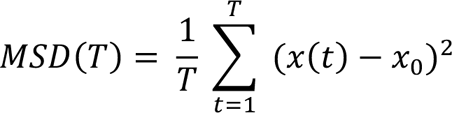

Equation 2:

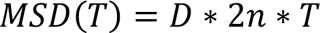

### Live cell CME visualization

To visualize adaptor protein recruitment in live cells, A549 WT cells were transfected with Epsin1-GFP (addgene, #22228) using Lipofectamine 2000 (LifeTechnologies). One day later, the cells were detached using trypsin and subsequently cultivated on top of immobilized viruses. Live-cell imaging of AP2-RFP and ClC-GFP was performed using gene-edited MDA-MB-231 cells^43^ (kindly provided by David Drubin). The cells were imaged using a Nikon Ti2 microscope equipped with a NikonApo TIRF 100x Oil DIC N2 NA 1.49 objective. The samples were excited using a 488 nm laser (Coherent) and emitted light was detected on a pco.edge 4.2 sCMOS camera (Excellitas). The cells were observed using TIRF illumination.

### F-actin labelling and imaging

For STORM imaging of F-actin, A549 WT cells were cultivated on top of immobilized viruses. At ∼80% confluency, the cells were fixed in 4% PFA for 20 min at RT. Samples were washed three times with PBS and blocked/permeabilized for 1h in blocking buffer (0.2% Triton, 0.2 % BSA in PBS). Afterwards, the cells were incubated in 0.56 µM phalloidin-A647 (Invitrogen, A22287) solution in PBS for 2h at RT in the dark. The samples were washed three times with PBS. STORM imaging was performed using an established photo-switching buffer made of 50 mM Tris with 10 mM NaCl, supplemented with 10% glucose, 50 mM 2-mercaptoethylamine, 0.5 mg ml^−1^ glucose oxidase and 40 µg ml^−1^ catalase. All chemicals were purchased from Sigma. For live-cell imaging of F-actin structures, A549 cells were transfected with LifeAct-GFP one day before detachment using Lipofectamine 2000 (LifeTechnologies) and subsequently cultivated on top of immobilized viruses. The samples were imaged as described above for live-cell CME visualization.

