## Supplementary Figures for "Single influenza A viruses induce nanoscale cellular reprogramming at the virus-cell interface"

[*] correspondence:

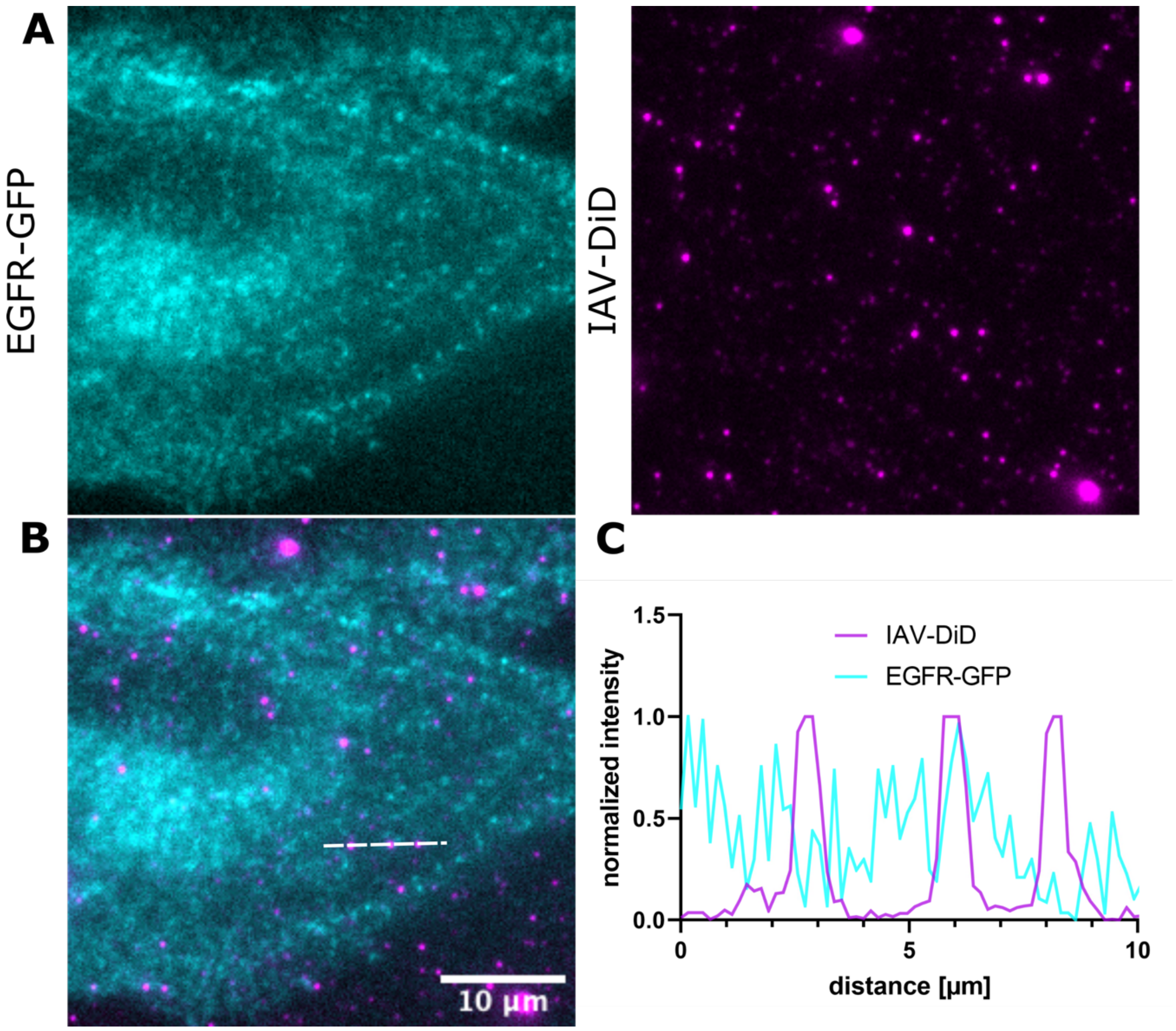


**Figure S1 | EGFR-GFP does not colocalize with IAV-DID after sialidase treatment**. A549 cells stably expressing EGFR-GFP (A) were cultivated in top of immobilized DID-labelled IAV PR8 (B). Before imaging, the cells were treated with 250 mU/ml sialidase for 1h. A line plot measurement (white line in B) shows no accumulation of EGFR-GFP at the virus-binding site (C).

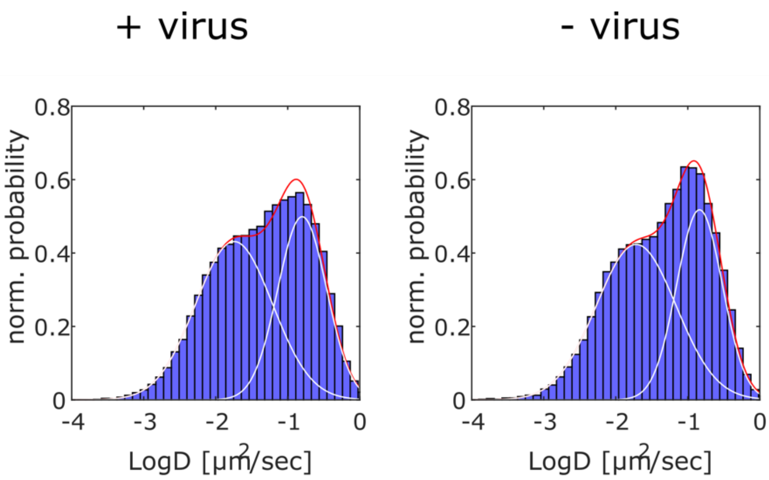


**Figure S2 | Histogram of EGFR diffusivity in cells cultivated with or without immobilized IAV**. A549 cells stably expressing EGFR-mEos3.2 were cultivated with or without immobilized IAV PR8. Histograms of the measured diffusion coefficients are shown. The scaled probability densities could be well fitted by two Gaussians (red curves). The white curves show the two subpopulations: left (right) for relatively immobile (mobile) tracks.

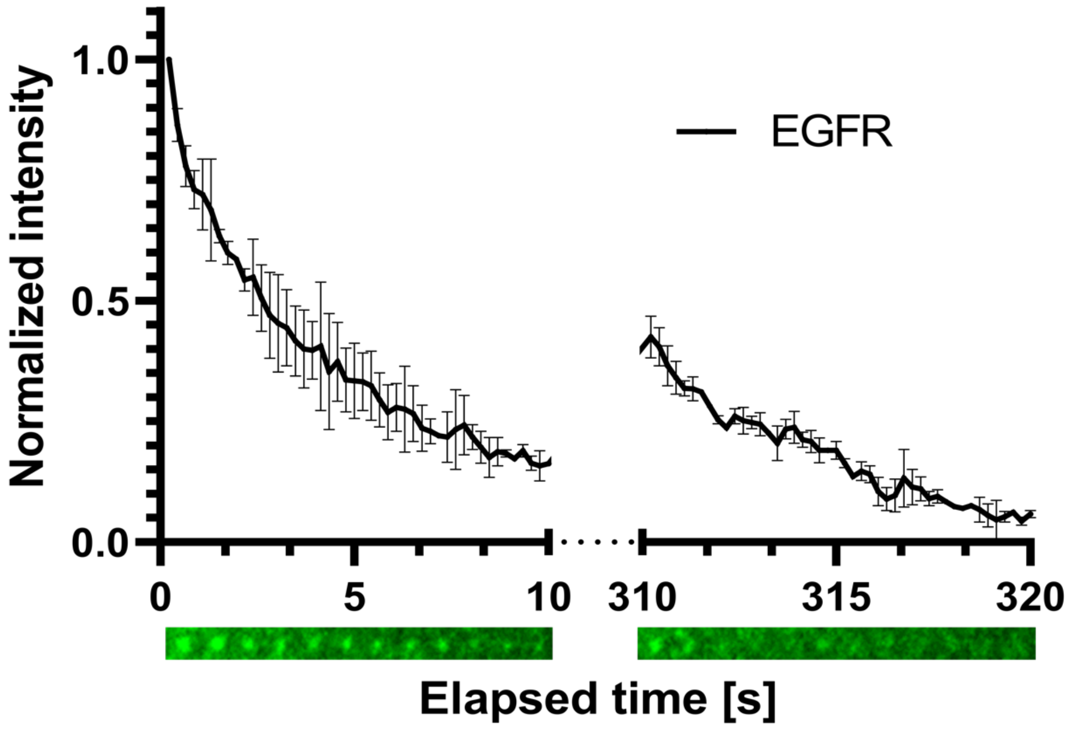


**Figure S3 | EGFR-GFP accumulated at the virus-binding site bleaches and recovers over time.** A549 cells stably expressing EGFR-GFP were cultivated in top of DiD-labelled IAV PR8. Interaction between IAV and EGFR led to an accumulation of EGFR-GFP at the virus-binding site (shown in **Figure 2A**). Continuous imaging of the same area led to photobleaching of the GFP signal (left side), which recovered after the excitation laser was switched of for 300 sec (right side).

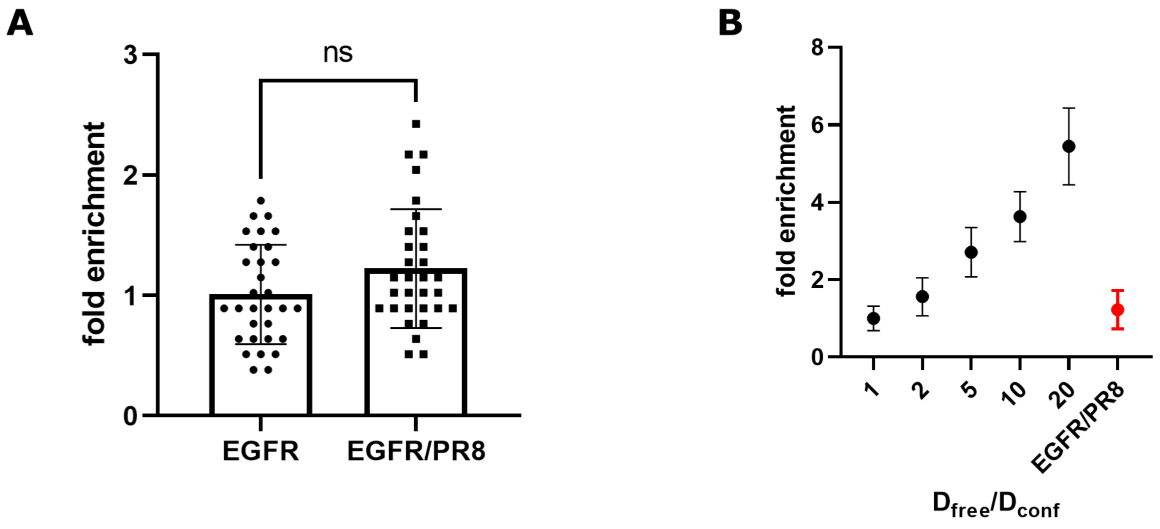


**Figure S4 | Virus-induced diffusion slowdown leads to local enrichment of receptor proteins.** To calculate if a slowdown of receptor proteins at the virus binding site leads to local protein enrichment, we simulated the 2D random diffusion of EGFR using a diffusion coefficient of $D_{EGFR}$ = 0.050 $\mu m^2/s$. We then added circular regions (r = 50 nm) to the simulation, where the proteins were slowed down according to our PALM measurements to $D_{EGFR-PR8}$ = 0.024 $\mu m^2/s$. We simulated 4000 time steps with 30 ms interval. At the end, we examined the number of receptors associated with the circular regions (i.e., viruses). (**A**) For the IAV PR8-mediated diffusion slowdown measured for EGFR, we find a moderate non-significant local enrichment as compared to freely diffusing receptors. (**B**) We also simulated stronger local slowdown as possible for other IAV-receptor combinations, which indeed leads to more pronounced local protein enrichment.

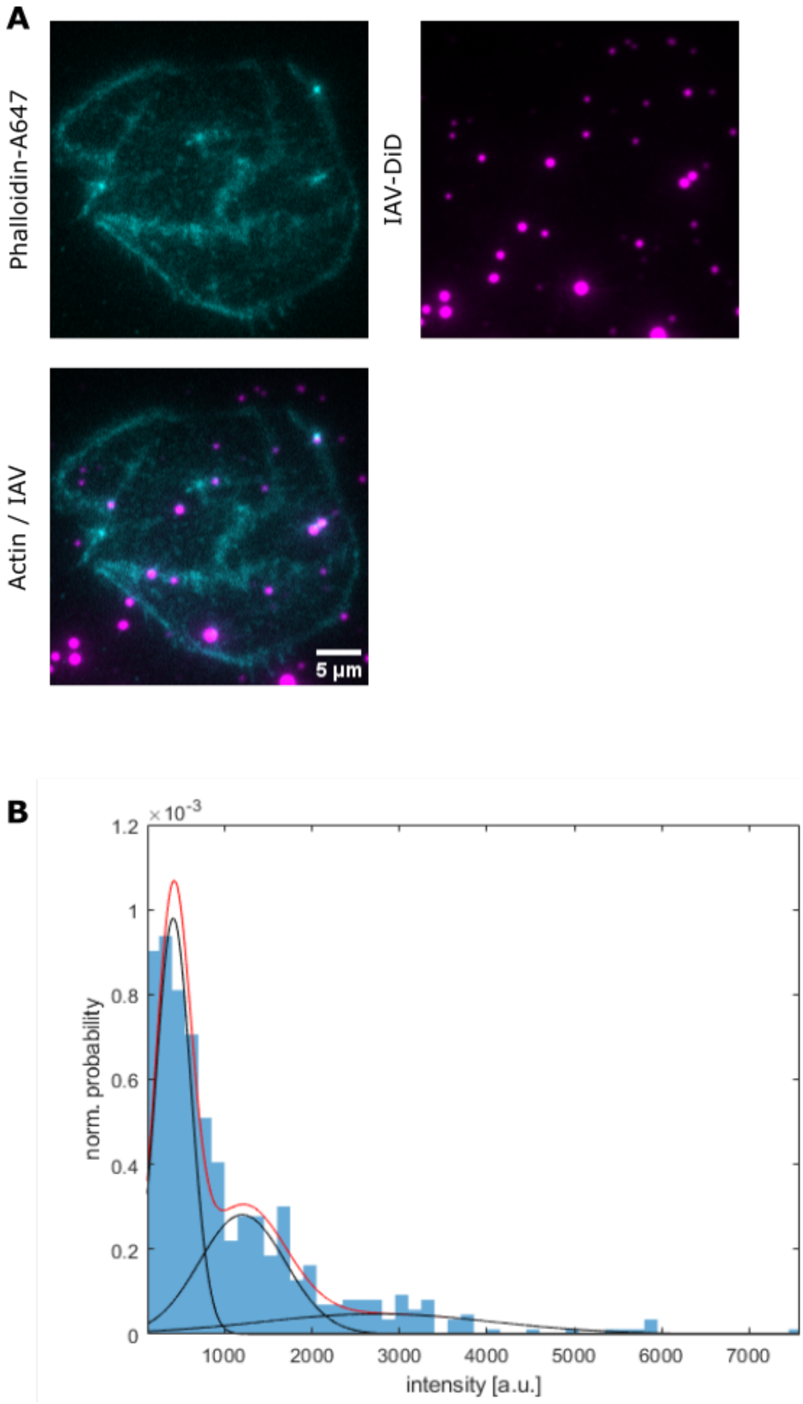


**Figure S5 | Immobilized IAV colocalize with phalloidin-Alexa647 in A549 cells**. A549 cells were cultivated on top of immobilized DiD-labelled IAV PR8. The cells were fixed and labelled using phalloidin conjugated with Alexa647 and imaged using TIRF illumination (**A**). For quantification, the viruses were detected automatically and the phalloidin-Alexa647 signal analyzed at the respective locations. The intensity distribution can be fitted using the sum of three Gaussians (B, red line). The three populations (B, black curves) were ascribed as 1) not below cells, 2) phalloidin low and 3) phalloidin high.

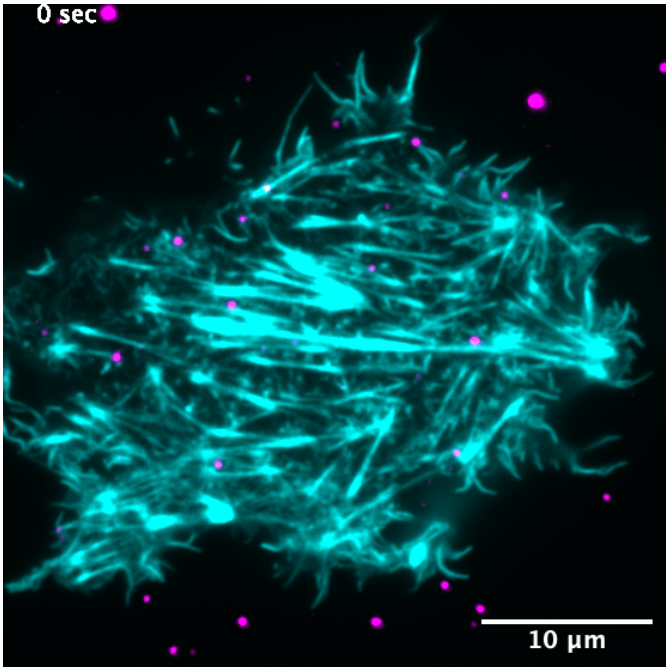


**Movie S1 | TIRF live-cell imaging of A549 cells expressing LifeAct-GFP cultivated on top of immobilized IAV**. A549 cells expressing LifeAct-GFP (cyan) were cultivated in top of immobilized DiD-labelled IAV PR8 and imaged by TIRF.
