## Supplementary figures and images for "Single influenza A viruses induce nanoscale cellular reprogramming at the virus-cell interface"

### Supplementary Video 1

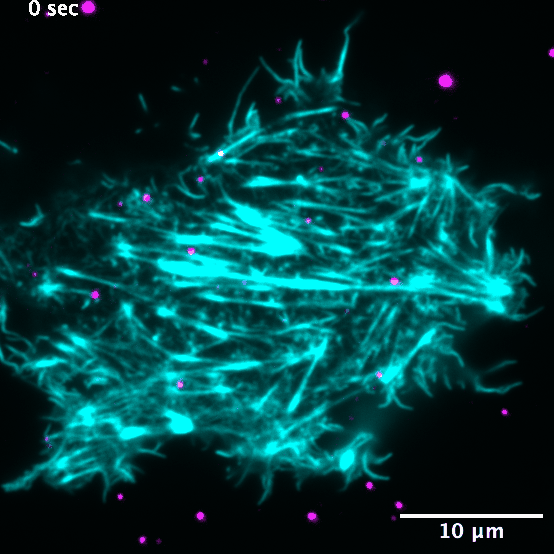
